# TrbA binds and locks a sliding clamp KorB to repress transcription on multi-drug resistance plasmids

**DOI:** 10.64898/2026.05.27.728120

**Authors:** Thomas C. McLean, Govind Chandra, Tung B.K. Le

## Abstract

Precise regulation of gene expression is essential for all living systems. Genes encoded on mobile genetic elements such as conjugative plasmids are also highly regulated to ensure stable inheritance, successful horizontal transfer, and adaptation to diverse bacterial hosts. The multi-drug resistance plasmid RK2 encodes an intricate regulatory network centred on the ParB-CTPase family protein KorB, which functions in both plasmid segregation and global gene regulation. We recently showed that KorB acts as a CTP-dependent DNA-sliding clamp that can be converted into a transcriptional repressor through direct interaction with the DNA-binding protein KorA. This clamp sliding-and-locking mechanism by KorAB enables highly effective transcriptional repression. Here, we investigate whether a third transcriptional regulator on the RK2 plasmid, TrbA, operates through a similar mechanism. Using a combination of structural prediction, biochemistry and *in vivo* transcriptional assays, we provide evidence that TrbA also directly interacts with KorB and functions as a clamp-locking factor. This KorB-TrbA interaction is mediated by a conserved aromatic interface that enables strong cooperative transcriptional repression. Disruption of this interface abolishes synergy, indicating that direct protein-protein interaction underpins KorB-TrbA cooperativity. Finally, bioinformatic analysis of a large plasmid database reveal that the tripartite KorB-KorA-TrbA system is present on numerous plasmids. Together, our findings establish TrbA as a *bona fid*e KorB co-repressor and demonstrate how a single sliding clamp protein can integrate multiple partners to evolve a complex transcriptional regulatory network.

**IMPORTANCE:** Precise regulation of gene expression ensures gene products are produced at the right time and in the right amounts. Recent works uncovered a new mechanism of bacterial gene regulation based on a clamp sliding and locking in a multi-drug resistance plasmid, RK2. KorB functions as a CTP-dependent DNA-sliding clamp capable of traveling over a long genomic distance. Sliding KorB is captured and locked in place by a partner protein, KorA, forming a stable complex at target promoters to repress transcription. Here, we show that another RK2 regulator, TrbA, also use this clamp sliding-locking mechanism, and identify an aromatic interface enabling TrbA-KorB-mediated transcriptional repression. Our findings show how a single sliding clamp integrates multiple partners to build a complex transcriptional regulatory network.

## INTRODUCTION

Precise regulation of gene expression is a fundamental requirement for all living organisms, enabling cells to coordinate complex biological processes by controlling when, where, and to what extent genes are transcribed. Despite their smaller genomic size, mobile genetic elements such as conjugative plasmids also need to dynamically regulate gene expression to ensure a stable vertical inheritance, successful horizontal transfer and adaptation to diverse host environments.

A striking example of complex transcriptional regulatory circuit is found in multi-drug resistance plasmids of incompatibility group P (IncP), which have been extensively studied due to their broad host range for both replication and conjugation (1). The archetypal RK2 plasmid has served as a model system for understanding the molecular basis of plasmid maintenance, gene regulation, and dissemination of antibiotic resistance (2, 3). Despite its small size, RK2 has a remarkably intricate regulatory network of four repressor proteins: KorA, KorB, KorC and TrbA (2, 4–7) (Fig. S1). KorB, a DNA-binding protein of the ParB CTPase family, plays a dual role in ensuring stable plasmid inheritance (a role analogous to chromosomal ParB in chromosome segregation) and in regulating gene expression for plasmid replication and conjugative transfer (4–10). We recently uncovered that KorB functions as a CTP-dependent DNA-sliding clamp, which can be converted into a potent transcriptional co-repressor by KorA (11). Analogous to chromosomal ParB (12–18), KorB loads onto its cognate DNA-binding site (*OB* or operator of KorB) while binding CTP, triggering engagement of the KorB homodimer N-terminal domains (NTD) and a conformational change that releases KorB from the *OB* site. This allows a now clamp-closed KorB to diffuse long-distances away from *OB* while remaining entrapped around the DNA (11). KorA bound at its DNA-binding site (*OA* or operator of KorA), through a direct protein-protein interaction, can capture the sliding KorB, thus turning the sliding clamp into a stable KorB-KorA-DNA complexes that are stationary near the target promoters. The KorAB complex on DNA creates a stable steric hindrance to prevent access of RNA polymerases to the target promoters, thereby repressing transcription (11). This KorA-KorB cooperative mechanism enhances transcriptional repression by ∼1000-fold compared to either protein acting alone, while at the same time enabling a mode of long-range transcriptional repression at promoters >1 kb from the *OB* site (11, 19–21). Transcriptional regulation by a DNA-sliding clamp, as observed for KorB-CTP, is analogous to of the activation of virulence genes in *Shigella flexneri* by a VirB-CTP sliding clamp (22–25) or the activation of T4-phage late genes by a phage-encoded PCNA-like Gp45 protein clamp (26, 27). The ability of KorA to trap sliding clamp KorB is functionally similar to eukaryotic CTCF and cohesin, respectively, where a DNA-binding protein CTCF restrains the translocation and loop extrusion by cohesin at specific DNA sites (28, 29). These examples suggest that “clamp sliding-and-locking” mechanisms may represent a general strategy for regulating gene expression in biology. In the context of the RK2 plasmid, previous works hinted at the possibility that the clamp sliding-and-locking mechanism might also operate with a third RK2 transcriptional regulator, TrbA.

TrbA (Fig. 1) regulates the expression of *trb* genes for pilus synthesis and mating pair formation (Fig. S1) and is therefore crucial for plasmid horizontal transmission (30–32). Similar to KorA, TrbA is also a small site-specific DNA-binding protein that recognizes an *OT* DNA sequence (operator of TrbA) (33) and exhibits weak transcriptional repressive activities on its own. Previous works have shown that TrbA requires KorB to achieve the full repression of *trb* genes (30, 34), but the molecular basis underlying this TrbA-KorB cooperation has not been fully resolved. Given the recent discovery of a clamp sliding-and-locking mechanism in KorAB, we hypothesized that TrbA and KorB might cooperate to heighten transcriptional repression in a similar manner.

**Fig. 1.**
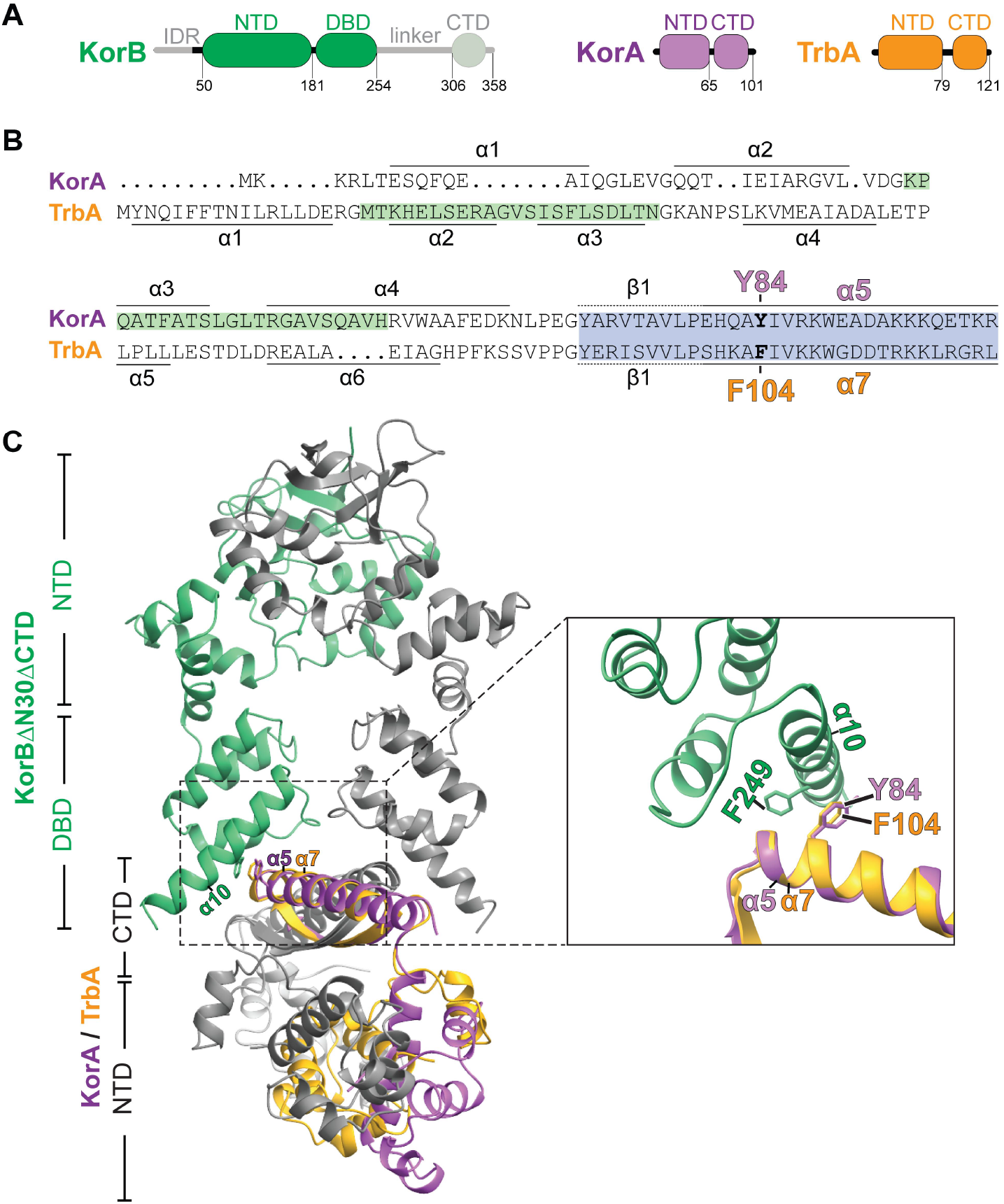
KorA and TrbA share structural homology at their C-terminal domains. **(A)** The domain architecture of KorB, KorA and TrbA. KorB consists of an N-terminal intrinsically disordered (IDR) IncC (ParA)-interacting peptide, a N-terminal CTPase domain (NTD), a central *OB* DNA-binding domain (DBD), a linker, and a C-terminal dimerization domain (CTD). The KorBΔN30ΔCTD variant used for crystallization lacks 30 N-terminal amino acids, linker domain and the CTD (faded green). KorA and TrbA both have N-terminal *OA* or *OT* DNA-binding domains (NTD) and C-terminal dimerization domains (CTD). **(B)** (left) The co-crystal structure of a KorBΔN30ΔCTD-KorA-DNA complex (PDB: 8QA9), the *OA* DNA in this structure is not shown here for simplicity. Overlayed on this structure is the AlphaFold2-predicted structure of homodimeric TrbA (yellow). (right) Helix α10 (green) of KorB interacts with helix α5 (purple) of KorA. Helix α7 (yellow) of TrbA aligns well with helix α5 of KorA, as does the critical aromatic residue KorA Y84 / TrbA F104. KorA Y84 interacts with KorB F249 through a π-π stacking interaction.

Here, using a combination of structural prediction, biochemical analysis, and *in vivo* gene expression assays, we provide evidence that TrbA is also a clamp-locking protein that directly interacts with the sliding clamp KorB. This interaction enables cooperative long-range transcriptional repression at specific *OT*-harbouring promoters on the RK2 plasmid. In particular, we identified a pair of aromatic residues on TrbA and KorB that mediate this interaction and are essential for cooperative transcriptional repression. Lastly, we bioinformatically survey the large and diverse collection of plasmids and suggest that the tripartite co-operative KorB-KorA-TrbA system appears widespread, extending beyond the archetypal RK2 plasmid. Altogether, our results show how a single sliding clamp KorB can integrate multiple interacting partners to widen the regulatory network.

## RESULTS & DISCUSSION

### KorA and TrbA share structural homology at their C-terminal domains

Cooperativity between KorB and KorA is mediated by a direct protein-protein interaction between the C-terminal domain of KorA (specifically helix α5) and helix α10 within the DNA-binding domain of KorB. It has been long been recognized that the C-terminal domains of KorA and TrbA are highly conserved, showing 76% similarity (55% identity) (Fig. 1A-B), while their N-terminal DNA-binding domains are more divergent (Fig. 1B) (19, 30, 31). This conservation suggests that TrbA might interact with KorB via a similar structural interface. To further explore this, we used AlphaFold2 (35) to predict the homodimeric structure of TrbA and the KorB-TrbA complex, and compared these models to the available X-ray crystallography structure of the KorB-KorA-DNA co-complex (PDB: 8QA9; Fig. 1C) (11). Consistent with sequence conservation, the C-terminal domains of KorA and TrbA closely aligned, while their N-terminal domains less so (Fig. 1B and Fig. S2). Structure alignment also suggests that, similar to the KorAB complex, the interaction interface between KorB and TrbA is formed between helix α10 of KorB, positioned just below the DNA-binding domain (KorB DBD; Fig. 1C), and the C-terminal helix α7 of TrbA, which forms part of the dimerization domain (TrbA CTD; Fig. 1C). Specifically, the F104 residue on helix α7 of TrbA align closely with the KorA Y84, a residue previously shown to mediate a key π-π stacking interaction with residue F249 on helix α10 of KorB (11) (Fig. 1C, right panel). Altogether, the high degree of sequence and predicted structural homology between TrbA and KorA, and the conservation of an aromatic residue at the equivalent interaction interface suggest that TrbA binds KorB via a similar mechanism as for KorA (11).

### TrbA and KorB directly interact

Using isothermal titration calorimetry (ITC) (Fig. 2A), we found that TrbA binds KorB *in vitro* with moderate affinity (K_D_ = 33.5 ± 1.86 µM), ∼tenfold weaker than KorA-KorB interaction (K_D_ = 4.2 ± 0.5 µM) (11). To determine if residue F104 of TrbA was critical for such interaction with KorB, we performed ITC using a purified TrbA (F104A) variant to find such alanine substitution eliminated the ability of TrbA to bind KorB (Fig. 2A).

**Fig. 2.**
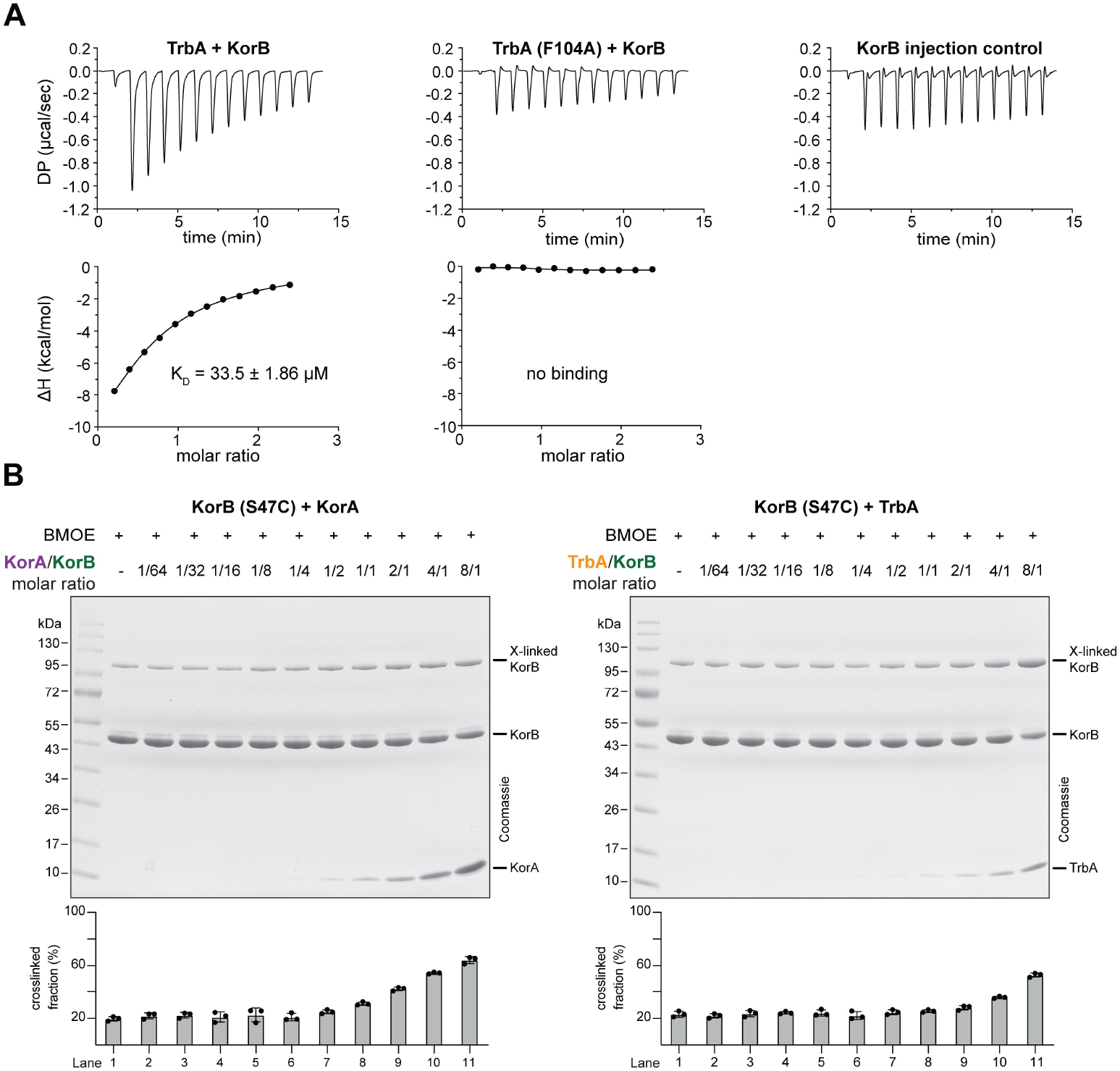
TrbA interacts with KorB *in vitro*. **(A)** Analysis of the interaction of TrbA (WT or F104A) with KorB (WT) by ITC. Experiment was duplicated and KorB-only injection control was also performed. **(B)** SDS-PAGE analysis of BMOE crosslinking products of 8 µM (dimer concentration) of KorB (S47C) ± increasing concentrations of: KorA (WT/Y84A) (left panel) or TrbA (WT/F104A) (right panel). Quantification of the crosslinked fraction is shown below each representative image. Experiments were performed in triplicate, and mean +/- SDs were presented.

### TrbA promotes the closing of the KorB clamp at its N-terminal domain

In the presence of CTP and *OB* DNA, KorB self-engages at its N-terminal domain (KorB NTD; Fig. 1A), a process called N-engagement, forming a closed clamp-like structure (11). KorA can induce KorB N-engagement even in the absence of CTP and *OB* DNA (11). Given the structural homology between KorA and TrbA and that TrbA directly interacts with KorB likely via the same protein-protein interface, we hypothesized that TrbA can similarly induce KorB N-engagement. We monitored N-engagement using cysteine-specific crosslinking of a purified KorB (S47C) variant with bismaleimidoethane (BMOE). Residue S47 at the NTD was substituted by cysteine on an otherwise cysteine-less KorB background; this substitution has previously been validated to not impact KorB activity while serving as an excellent proxy to detect N-engagement via crosslinking assay (11).

In the presence of BMOE, KorB (S47C) alone crosslinked at ∼20% (Fig. 2B, lane 1). Addition of KorA (Fig. 2B, left) or TrbA (Fig. 2B, right) across increasing molar ratios relative to KorB, from 1:512 (lane 2) to equimolar (lane 11), increased KorB (S47C) crosslinking efficiency to ∼60% and ∼40%, respectively. The results indicate that, like KorA, TrbA can also promote KorB N-engagement. We noted that crosslinking efficiency above the basal level (of ∼20%) was detected as early as when KorA was added at only 1:4 molar ratio (lane 8), but not for TrbA at the equivalent molar ratio. Since KorB N-engagement likely requires direct binding of KorA/TrbA to KorB, the results here are consistent with KorB binding TrbA with a weaker affinity than to KorA, as observed by ITC (Fig. 2A).

Next, we investigated whether the conserved residue F104 in TrbA is required for TrbA-stimulated KorB N-engagement (Fig. 3A). As previously observed, the basal level of KorB (S47C) crosslinking in the presence of BMOE alone was ∼20% (Fig. 3B, lane 1). No crosslinking of KorB (S47C), KorA or TrbA was observed in the absence of BMOE (Fig. 3A, lanes 2 and 3; KorA and TrbA are cysteine-less). Addition of equimolar TrbA or KorA (positive control) increased crosslinking efficiency to ∼55% (lanes 4 and 5). The crosslinking efficiency of KorB (S47C) increased further to ∼70% in the presence of both CTP and cognate *OB*-DNA (lane 6) and was unchanged by the subsequent addition of TrbA or KorA (lanes 7 and 8), suggesting that KorB NTD engagement is maximal under these conditions. Consistent with a previous report, KorA (Y84A), defective in KorB binding, failed to increase crosslinking above basal levels (lane 9) (11). Similarly, the TrbA (F104A) variant was unable to promote N-engagement beyond ∼20% (Fig. 3A, lane 10). Together with the ITC data (Fig. 2A), these results indicate that TrbA promotes KorB N-engagement likely through a direct interaction dependent on the conserved aromatic residue F104.

**Fig. 3.**
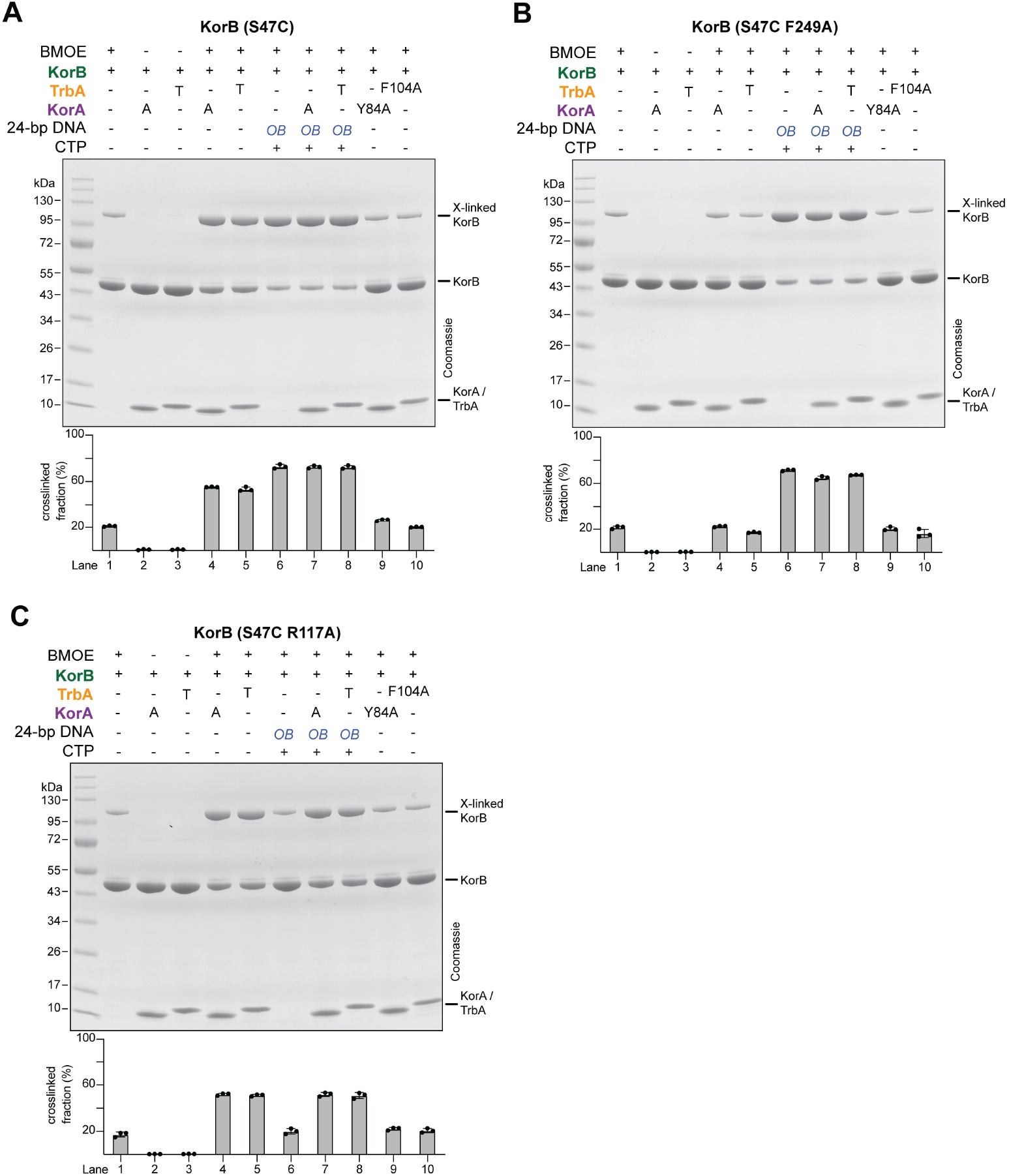
TrbA promotes the N-engagement of KorB *in vitro*. SDS-PAGE analysis of BMOE crosslinking products of 8 µM (dimer concentration) of **(A)** KorB (S47C), **(B)** KorB (S47C F249A), or **(C)** KorB (S47C R117A) ± 8 µM (dimer concentration) of TrbA (WT/F104A) or KorA (WT/Y84A) ± 1 µM *OB* DNA ± 1 mM CTP. Quantification of the crosslinked fraction is shown below each representative image. Experiments were performed in triplicate, and mean +/- SDs were presented.

To probe the KorB side of this interaction, we introduced the F249A substitution in KorB, previously shown to disrupt KorA binding (11). KorB (S47C F249A) retained its intrinsic capacity to undergo N-engagement, with crosslinking efficiency reaching ∼70% in the presence of CTP and cognate *OB*-DNA (lane 6) i.e. comparable to KorB (S47C) (Fig. 3A, lane 6). However, neither KorA or TrbA enhanced the crosslinking of KorB (S47C F149A) in any tested condition (Fig. 3B, lanes 4, 5, 7 and 8). Not to our surprise, neither KorA (Y84A) and TrbA (F104A) were able to further increase KorB (S47C F249A) crosslinking either (Fig. 3B, lanes 9 and 10). These results are consistent with our previous report that F249 on KorB and Y84/F104 on its binding partners KorA/TrbA are required for protein-protein interaction and KorB N-engagement (11).

The KorB (R117A) variant, unable to bind CTP, is impaired in N-engagement (11). Having defined the KorB-TrbA interaction interface, we next asked whether TrbA could promote N-engagement in KorB (R117A) independently of CTP binding (Fig. 3C). Again, the basal crosslinking level of KorB (S47C R117A) in the presence of BMOE alone was ∼20% (Fig. 3C, lane 1). KorB (S47C R117A) alone was unresponsive to the addition of CTP and *OB*-DNA, with the level of crosslinking remaining at ∼20% (Fig. 3C, lane 6). Nevertheless, addition of equimolar KorA or TrbA increased crosslinking efficiency to ∼60% for both variants (lanes 4 and 5), indicating that TrbA, like KorA, can promote KorB N-engagement independently of CTP binding. Again, alanine substitution of Y84 and F104 in KorA or TrbA reduced crosslinking efficiency back to basal levels (Fig. 3C, lanes 9 and 10). Altogether, our results indicate that TrbA, like KorA, promotes KorB N-engagement independently of CTP and cognate *OB* DNA, and that this clamp-closing activity is dependent on a conserved pair of aromatic residues from both proteins.

### TrbA-KorB interaction mediates cooperative transcriptional repression

TrbA has previously been shown to cooperate with KorB to strongly repress gene transcription on the RK2 plasmid (19, 30, 33), but whether this cooperation depends on a direct physical interaction between the two proteins has remained untested. To investigate, we constructed two promoter-*xylE* transcriptional fusion reporters and measured catechol dioxygenase activities in the presence or absence of KorB (WT/variants) and/or TrbA (WT/variants) *in vivo* (Fig. 4A). In these reporters, the KorB-binding site (*OB*) was positioned either 4 bp (*OB*-proximal) or 1.6 kb (*OB*-distal) upstream of the core promoter −35 element from the RK2 gene *trbB*, while the TrbA-binding site (*OT*) overlaps with the −10 element, mimicking KorB-TrbA-regulated promoters natively found on the RK2 plasmid (Fig. 4A).

**Fig. 4.**
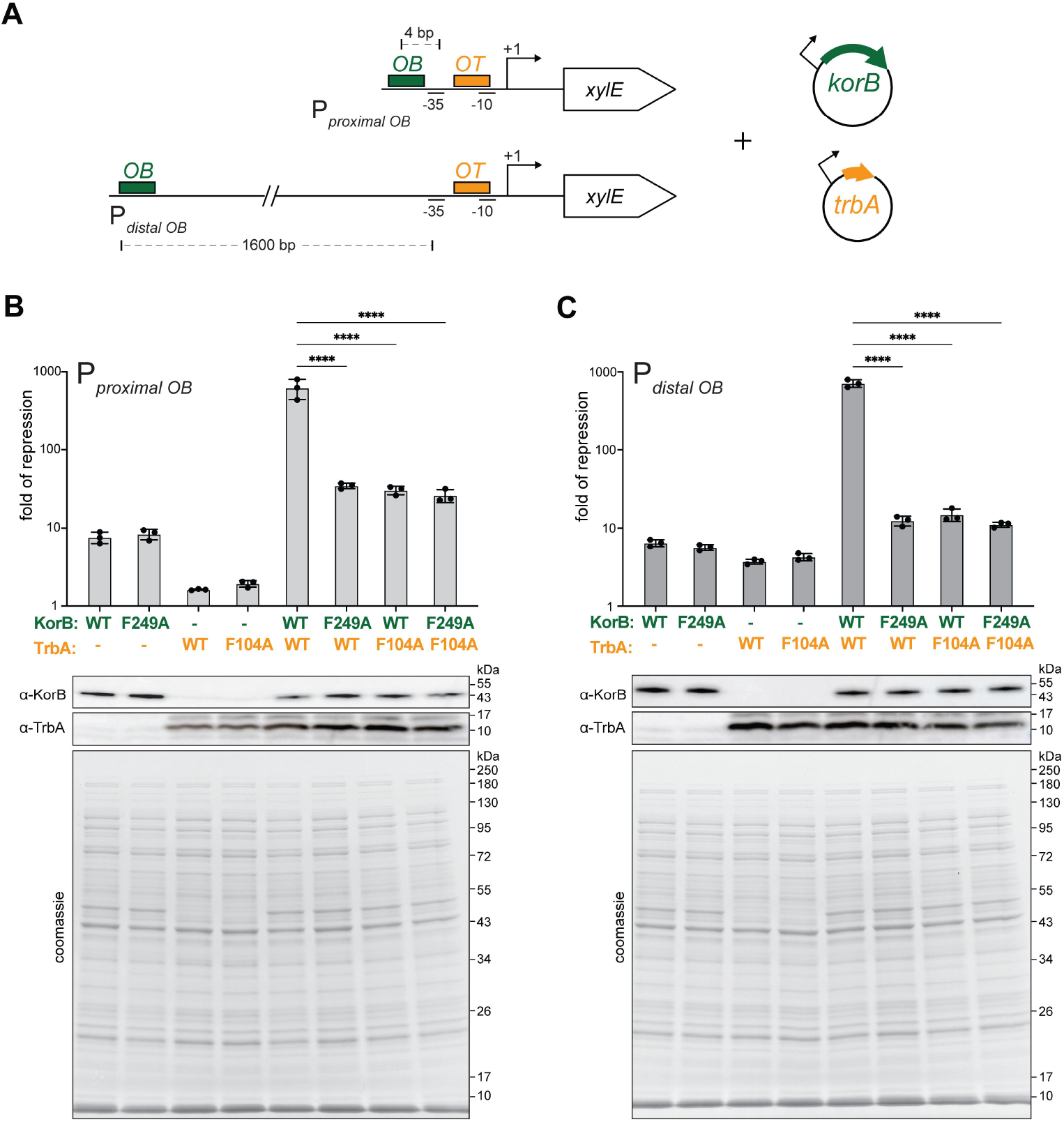
TrbA cooperates with KorB to repress transcription *in vivo*. **(A)** Schematic diagrams of promoter-*xylE* reporter constructs. **(B)** (left panel) Fold of repression for proximal promoter reporter. (right panel) Fold of repression for distal promoter reporter. Fold of repression is a ratio of XylE activities from cells co-harboring a reporter plasmid and KorB/TrbA expressing plasmids to that of cells co-harboring a reporter plasmid and an empty plasmid (KorB/TrbA minus control). Immunoblots of α-TrbA and α-KorB from lysates of cells used in the same experiments are also shown. Experiments were performed in triplicate and mean ± SDs were presented. See also supplementary Fig. S3 for absolute values of XylE activities.

Consistent with previously reports (19, 30, 33), KorB alone produced weak repression of both the proximal (∼8 fold; Fig. 4B, lane 1) and distal (∼6 fold; Fig. 4C, lane 1) promoters (see also Fig. S1 for absolute values from promoter-*xylE* reporter assays). TrbA alone had an even weaker repression of ∼2 fold (Fig. 4B, lane 3) and ∼4 fold (Fig. 4C, lane 3) for the proximal and distal promoters, respectively. Notably, co-expression of KorB and TrbA, however, increased transcriptional repression markedly to over 600-fold for both promoters (Fig. 4B and C, lane 5). To quantify the synergistic contribution from both proteins, we calculated a cooperation coefficient by dividing the observed repression level by the expected repression as if the two proteins acted independently (i.e. KorB WT alone * TrbA WT alone). This yielded co-operation coefficients of 51 and 30 for the proximal and distal promoters respectively, indicative of highly synergistic activity.

We next repeated the experiments using the interaction-deficient variants KorB (F249A) and TrbA (Y84A). Immunoblots using α-KorB and α-TrbA confirmed that all KorB and TrbA variants were produced at the similar level to wild-type. We observed that, when expressed individually, each variant repressed transcription similarly to its wild-type counterpart (Fig. 4B and C, lanes 2 and 4). However, co-expression of these variants failed to produce cooperative repression observed with wild-type proteins, with co-operation coefficients reducing to 2 at the proximal promoter (Fig. 4B, lanes 6-8) and 0.5 at the distal promoter (Fig. 4C, lanes 6-8) i.e. a complete loss of cooperativity. Altogether, our results show that direct interaction between KorB and TrbA, mediated through the key aromatic residues, is essential for cooperative transcriptional repression for both *OB*-proximal and long-range *OB*-distal promoters.

### The tripartite KorB-KorA-TrbA system is present on many plasmids

To assess the distribution of this cooperative repression system, we bioinformatically searched the PLSDB plasmid database (∼70,000 entries) (36) for KorB, KorA, and TrbA homologs. Protein BLAST searches were performed using full-length RK2 KorA and TrbA, and residues 39-149 of KorB that spans the conserved NTD and DNA-binding domain, as queries. In total, we identified 5,871 KorB, 1,385 KorA and 1,036 TrbA homologs across 6,161 unique plasmid accessions.

To evaluate the potential for cooperative interactions, sequence alignments were constructed around the key aromatic residues required for KorB-KorA/TrbA interaction i.e. F249 in RK2 KorB, Y84 in RK2 KorA, and F104 in RK2 TrbA. Homologues were subsequently classified based on the presence or absence of aromatic residues (phenylalanine, tyrosine, or tryptophan) at the equivalent position. Of the 6,161 accessions, the majority i.e. 5,566 (90.3% of all plasmid accessions) contained one, 267 (4.3%) contained two, and 328 (5.3%) contained all three: KorB, KorA and TrbA homologues (Fig. 5). Of the single-protein cases, KorB was the most prevalent (∼80%), consistent with the core role of the ParB-like KorB in vertical plasmid segregation, whereas KorA and TrbA are rarer (found without KorB in 13.6% and 6.6% of cases, respectively).

**Fig. 5.**
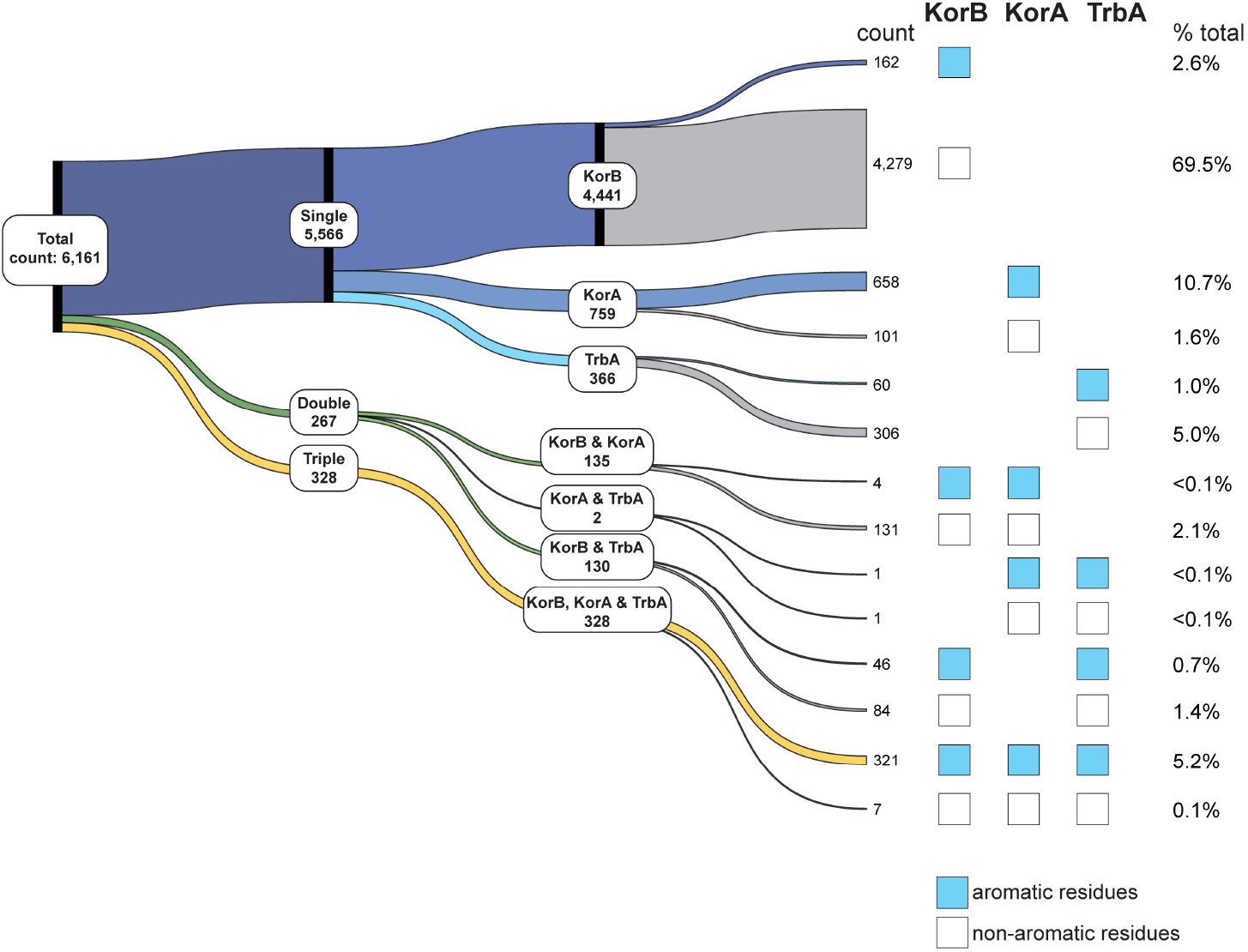
The tripartite KorB-KorA-TrbA system is present on many plasmids. (left panel) Sankey diagram depicting the presence of genes encoding *korB, trbA* and *korA* homologs from plasmid accessions in PLSDB plasmid database. Single indicates the presence of only one protein i.e. only KorB, only KorA, or only TrbA homolog. Double indicates the presence of two proteins. Triple indicates the presence of all three KorB, KorA, and TrbA homologs on the same plasmid. (right panel) Analysis of the conservation of aromatic residues at the KorB-KorA/TrbA interface.

Analysis of the conservation of aromatic residues at the KorB-KorA/TrbA interface revealed distinct patterns. KorB and TrbA homologs found in isolation predominantly lack aromatic residues at this interface, while 86.7% of KorA homologs found alone have aromatic residues at this position. Among the 135 plasmids containing both KorB and KorA, only four plasmids (∼3%) carried aromatic residues on both proteins, suggesting a potential functional cooperative system is rare in this subset. A similar analysis of the 130 plasmids containing both KorB and TrbA revealed that 46 plasmids (35.4%) encoded aromatic-positive variants of both proteins, suggesting a higher likelihood of KorB-TrbA cooperation in this subset. Most notably, of the 328 plasmids encoding all three proteins, the majority, i.e. 321 plasmids (97.9%), contained aromatic-positive variants of KorB, KorA and TrbA, suggestive of a fully intact tripartite co-operative system analogous to that on RK2 (Fig. 5).

Taken together, this bioinformatic survey shows that homologs of KorB, KorA and/or TrbA are well presented in ∼10% of all plasmids in the PLSDB, with KorB being the most widespread, likely reflecting its essential role in plasmid segregation. Whilst the complete tripartite KorB-KorA-TrbA system presents in only ∼5.3% of plasmids, the near-universal conservation of the aromatic residues at the protein-protein interaction interface in these cases is suggestive that cooperation is under selective pressure. It should be noted that the PLSDB database represents only a fraction of global plasmid diversity, prioritizing curation and quality over coverage compared to larger alternative repositories (19). The true prevalence of these co-operative systems, and whether there exists KorB partners beyond KorA and TrbA, across the broader plasmid metagenome therefore remains unclear.

In summary, findings in this work establish TrbA as a *bona fide* KorB co-repressor that operates through the same conserved aromatic interface as KorA (Fig. 6). By analogy with KorA, we speculate that TrbA bound to an *OT* DNA site near the target promoter would capture KorB as it slides from the distal *OB* site. Upon interaction, TrbA locks KorB in a closed-clamp conformation, potentially preventing its release from DNA. Thus, this TrbA-KorB complex would then form a stationary complex at the target promoter, likely occluding RNA polymerases from accessing the core promoter site to initiate transcription (Fig. 6). Because TrbA recognizes a DNA sequence distinct from that of KorA, KorB’s ability to exploit the same interaction interface with both KorA & TrbA represents an elegant strategy for integrating multiple regulatory partners to controls distinct set of genes and build a more complex transcriptional network. These insights deepen our understanding of the multilayered transcriptional regulatory network encoded by RK2 and highlight the conserved aromatic interface as a potential target for disrupting plasmid maintenance in multi-drug resistance bacteria.

**Fig. 6.**
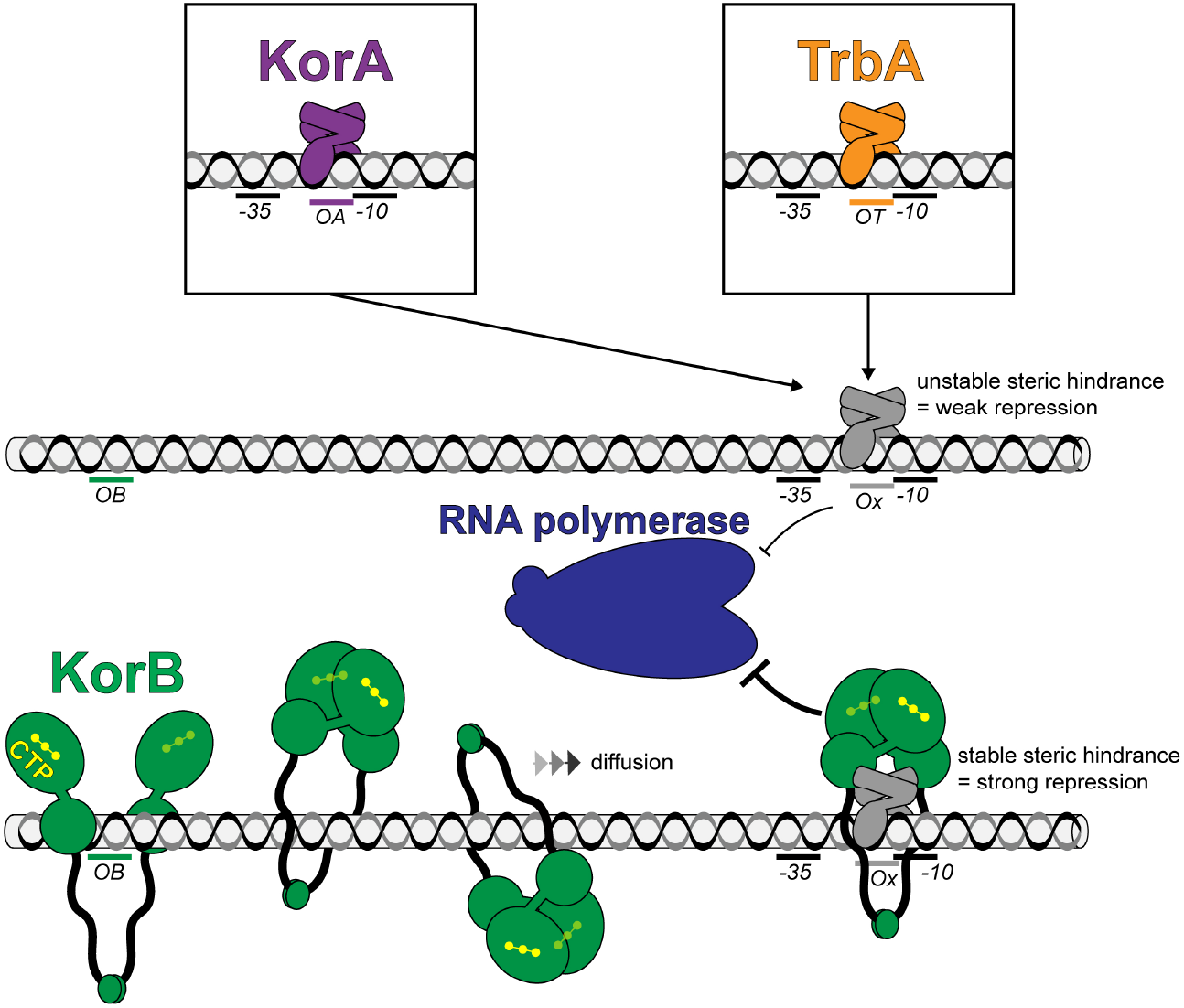
A proposed model for the cooperativity between TrbA/KorA and KorB for long-range transcription repression. On the RK2 plasmid the binding site for TrbA (*OT*) or KorA (*OA*) are typically found overlapping the −10 element of the core promoter site. The KorB binding site (*OB*) can be positioned kb away from the core promoter. CTP-binding triggers KorB to clamp around DNA (N-engagement) and slide via 1D diffusion for kb distances along the DNA. Through this movement KorB can encounter and bind DNA-bound TrbA/KorA at the core promoter site. This produces a stable co-complex that occludes RNA polymerases from accessing the core promoter site.

### Data and code availability

All images and source data presented in figures are available in Mendeley (DOI: 10.17632/6v6gr2875g.1). This paper does not report original code.

### Declaration of interests

The authors declare no competing interests.

## MATERIALS AND METHODS

### Strains, media, and growth conditions

*E. coli* strains were grown in lysogeny broth (LB) medium. When appropriate, the media was supplemented with antibiotics at the following concentrations (liquid/solid [μg mL^-1^]): carbenicillin (50/100), chloramphenicol (20/30), kanamycin (30/50) and streptomycin (50/50).

### Plasmid and strain construction

Plasmids and strains used or generated in this study are listed in Supplementary Table S1.

For plasmid construction, a double-stranded (ds) DNA fragment containing a desired sequence was chemically synthesized (gBlocks, IDT). The target plasmid was double-digested with restriction enzymes and gel-purified. A 10 μL reaction mixture was created with 5 μL 2x Gibson master mix (NEB) and 5 μL of combined equimolar concentration of purified plasmid backbone and gBlock(s).

This mixture was incubated at 50◦C for 60 min. Gibson assembly was possible owing to shared sequence similarity between the digested plasmid backbone and the gBlock fragment(s). All resulting plasmids were verified by Sanger sequencing (Genewiz) or whole-plasmid sequencing (Plasmidsaurus).

#### Construction of pET21b::trbA-strep (WT and mutants)

DNA fragments containing WT/mutated *trbA* genes (*trbA*\*) were chemically synthesized with a C-terminal Strep-tag sequence (gBlocks, IDT). The NdeI-HindIII-cut pET21b plasmid backbone and *trbA*\* gBlock fragments were assembled using a 2x Gibson master mix (NEB). Gibson assembly was possible owing to a 37 bp sequence shared between the NdeI-HindIII-cut pET21b backbone and the gBlocks fragment.

#### Construction of pDM1.2::trbA (mutants)

DNA fragments containing mutated *trbA* genes (*trbA\**) were chemically synthesized (gBlocks, IDT). The EcoRI-SalI-cut pDM2.1 plasmid backbone and *trbA\** gBlocks fragments were assembled using a 2x Gibson master mix (NEB). Gibson assembly was possible owing to a 38 bp sequence shared between the EcoRI-SalI-cut pDM2.1 backbone and the gBlocks fragment.

#### Construction of E. coli DH5α and BL21 pLysS strains containing the desired combinations of plasmids

Plasmids were introduced/co-introduced into *E. coli* DH5α or *E. coli* BL21 pLysS via heat shock transformation (42°C, 30 s) in the required combinations.

### Protein overexpression and purification

#### KorB (WT and mutants)

Protein preparation of His_6_-tagged KorB (WT and mutants) for ITC was performed as follows. C-terminally His-tagged KorB (WT and mutants) were expressed from the plasmid pET21b in *E. coli* Rosetta (BL21 DE3) pLysS competent cells (Merck). 80 mL of overnight culture was used to inoculate 4 L of LB with selective antibiotics. Cultures were incubated at 37°C with shaking at 200 rpm until OD_600_ ∼0.6. Cultures were cooled for 1 hr at 4°C before isopropyl-β-D-1-thiogalactopyranoside (IPTG) was added to a final concentration of 0.5 mM. The cultures were incubated overnight at 16°C with shaking at 200 rpm before cells were pelleted by centrifugation. Cell pellets were resuspended in buffer A (100 mM Tris-HCl, 300 mM NaCl, 10 mM imidazole, 5% v/v glycerol, pH 8.0) with 5 mg lysozyme (Merck) and a cOmplete EDTA-free protease inhibitor cocktail tablet (Merck) at room temperature for 30 min with gentle rotation. Cells were lysed on ice with 10 cycles of sonication: 15 s on / 15 s off at an amplitude of 20 microns, pelleted at 16,000 rpm for 35 mins at 4°C and the supernatant filtered through a 0.22 µm sterile filter (Sartorius). Clarified lysate was loaded onto two 1 mL HisTrap HP columns (Cytiva) pre-equilibrated with buffer A. Protein was eluted from the column using an increasing gradient of imidazole (10-500 mM) in the same buffer. Desired protein fractions were pooled and loaded onto a preparative-grade HiLoad 16/600 Superdex 200pg gel filtration column (GE Healthcare) pre-equilibrated with elution buffer (100 mM Tris-HCl, 300 mM NaCl, 5% v/v glycerol, pH 8.0). Desired fractions were identified and analysed for purity via SDS-PAGE before being pooled. Aliquots were flash-frozen in liquid nitrogen and stored at −80°C.

Protein preparations for BMOE crosslinking assays were purified using a one-step Ni-affinity column with all buffers adjusted to pH 7.4 for optimal crosslinking. Purified proteins were subsequently desalted using a PD-10 column (Merck) before being concentrated using an Amicon Ultra-4 10 kDa cut-off spin column (Merck). Final protein samples were aliquoted and stored at −80°C in storage buffer (100 mM Tris-HCl, 300 mM NaCl, 5% v/v glycerol, 1 mM Tris(2-carboxyethyl) phosphine, pH 7.4).

#### KorA (WT and mutants)

Protein preparation of His_6_-tagged KorA (WT and mutants) for BMOE crosslinking was performed as above for KorB (WT and mutants) with one exception. Following elution from the HisTrap HP columns (Cytiva) the protein fractions were pooled and loaded onto a preparative-grade HiLoad 16/600 Superdex 75pg gel filtration column (GE Healthcare) pre-equilibrated with elution buffer (100 mM Tris-HCl, 300 mM NaCl, 5% v/v glycerol, pH 8.0).

#### TrbA (WT and mutants)

Strep-tagged TrbA (WT and mutants) for BMOE crosslinking and ITC were purified as follows. C-terminally Strep-tagged TrbA (WT and mutants) were expressed from the plasmid pET21b in *E. coli* Rosetta (BL21 DE3) pLysS competent cells (Merck). 120 mL of overnight culture was used to inoculate 6 L of LB with selective antibiotics. Cultures were incubated at 37°C with shaking at 200 rpm until OD_600_ ∼0.6. Cultures were cooled for 1 hr at 4°C before isopropyl-β-D-1-thiogalactopyranoside (IPTG) was added to a final concentration of 0.5 mM. The cultures were incubated overnight at 16°C with shaking at 200 rpm before cells were pelleted by centrifugation. Cell pellets were resuspended in buffer A (100 mM Tris-HCl, 300 mM NaCl, 5% v/v glycerol, pH 8.0) with 5 mg lysozyme (Merck) and a cOmplete EDTA-free protease inhibitor cocktail tablet (Merck) at room temperature for 30 min with gentle rotation. Cells were lysed on ice with 10 cycles of sonication: 15 s on / 15 s off at an amplitude of 20 microns, pelleted at 16,000 rpm for 35 mins at 4°C and the supernatant filtered through a 0.22 µm sterile filter (Sartorius). 15 mL of clarified lysate was loaded onto a 1 mL StrepTrap XT column (Cytiva) pre-equilibrated with buffer A. Protein was eluted from the column using 5 mL buffer B (100 mM Tris-HCl, 300 mM NaCl, 100 mM biotin, 5% v/v glycerol, pH 8.0) in 0.25 mL fractions into 2 mL wells pre-loaded with 1 mL of buffer A. Desired protein fractions were pooled and loaded onto a preparative-grade HiLoad 16/600 Superdex 75pg gel filtration column (GE Healthcare) pre-equilibrated with elution buffer (100 mM Tris-HCl, 300 mM NaCl, 5% v/v glycerol, pH 8.0). Desired fractions were identified and analysed for purity via SDS-PAGE before being pooled. Aliquots were flash-frozen in liquid nitrogen and stored at −80°C.

Both biological (new sample preparations from a stock aliquot) and technical (same sample preparation) replicates were performed for assays in this study. Protein concentrations were determined by Bradford assay and reported as concentrations of KorA, TrbA or KorB dimers.

### DNA preparation for *in vitro* crosslinking

Palindromic single-stranded DNA oligonucleotides (*OB*: GGGAT<u>ATTTTAGCGGCTAAAAG</u>GA) were resuspended to 100 µM in 1 mM Tris-HCl, 5 mM NaCl, pH 8.0 buffer and were heated at 98°C for 5 min before being left to cool down to room temperature overnight to form 50 µM double-stranded DNA. The core sequence of *OB* is underlined.

### *In vitro* crosslinking assay using a sulfhydryl-to-sulfhydryl crosslinker BMOE

A 50 µL mixture of 8 µM KorB WT or mutants ± 1 mM NTP ± 0.5 µM 24-bp dsDNA containing *OB*-DNA was assembled in a reaction buffer (10 mM Tris-HCl pH 7.4, 100 mM NaCl, and 1 mM MgCl_2_) and incubated for 5 min at room temperature. BMOE was added to a final concentration of 1 mM, and the reaction was quickly mixed by three pulses of vortexing. The reaction was then immediately quenched through the addition of SDS-PAGE sample buffer containing 23 mM β-mercaptoethanol. Samples were heated to 80°C for 5 min before being loaded on 4-20% Novex WedgeWell Tris-Glycine gels (Thermo Fisher Scientific). Protein bands were stained with an InstantBlue Coomassie protein stain (Abcam), and band intensity was quantified using ImageJ. The results were analysed in Excel and plotted using GraphPad Prism 10.

### *xylE* reporter gene assays

Reporter gene assays were carried out as in McLean *et al* (2025) (11). In short, *E. coli* DH5α cells containing relevant expression plasmids were grown to a logarithmic phase from a 1:100 dilution of overnight culture. Induction of KorB WT/mutant expression was achieved with 0.2% arabinose. Induction of TrbA WT/F104A required no additional IPTG. 50 mL of culture was pelleted and resuspended in 500 μL resuspension buffer (0.1 M sodium phosphate buffer pH 7.4, 10% v/v acetone). From this point onwards samples were kept on ice. Cells were disrupted using sonication at 10 microns for 10 sec and subsequently pelleted. The supernatant was transferred to a fresh microcentrifuge tube and assayed for catechol 2,3-oxygenase activity. Samples were diluted 1:10 in reaction buffer (0.1 M sodium phosphate buffer pH 7.4, 200 μM catechol) and incubated at room temperature for a minute before the absorbance at 374 nm was determined using a BioMate 3 spectrophotometer (Thermo Fisher Scientific). Protein concentration, determined using Bradford assay, was used to normalize the samples. The results were analysed in Excel and plotted using GraphPad Prism 10.

### Immunoblot analysis

For immunoblots, 50 ng (α-KorB immunoblots) or 2 µg (α-TrbA immunoblots) of total protein lysate was resuspended in 1X SDS-PAGE sample buffer and heated to 95°C for 10 min before being loaded onto each well. Denatured samples were run on 12% Novex WedgeWell gels (for α-KorB immunoblots) or 4-20% Novex WedgeWell gels (for α-TrbA immunoblots) at 150 V for 55 mins. Resolved proteins were transferred to PVDF membranes using the Trans-Blot Turbo Transfer System (BioRad) and the membrane subsequently transferred to an iBind Western Device (Thermo Fisher Scientific). The membrane was processed following manufacturer’s instructions with a 1:10,000 dilution of α-KorB primary antibody (Cambridge Research Biochemicals) or with 1:4,000 dilution of α-TrbA (12) followed by a 1:10,000 dilution of goat α-rabbit HRP-conjugated secondary antibody (Abcam). Blots were imaged after incubation with SuperSignal West PICO PLUS Chemiluminescent Substrate (Thermo Fisher Scientific) using an Amersham Imager 600 (GE HealthCare). Loading controls of denatured 200 ng total protein lysate were run on 12% Novex WedgeWell gels (Thermo Fisher Scientific) at 150 V for 55 mins and stained with InstantBlue Coomassie protein stain (Abcam).

### Isothermal titration calorimetry (ITC)

All ITC experiments were recorded using a MicroCal PEAQ ITC instrument (Malvern Panalytical). Experiments were performed at 25°C. All components for binding experiments were in 100 mM Tris-HCl, 300 mM NaCl, 5% v/v glycerol, pH 8.0 buffer. For each ITC run, the calorimetric cell was filled with 40 μM dimer concentration of TrbA (WT or mutant) and a single injection of 0.4 μL of 500 μM KorB (WT) was performed first, followed by 12 injections of 3 μL each. Injections were carried out at 60 s intervals with a stirring speed of 750 rpm. Controls of KorB (WT) into buffer and buffer into TrbA (WT or mutant) were performed with no binding signal observed. The raw titration data were integrated using the binding control data and fitted to a one-site binding model using the built-in software of the MicroCal PEAQ ITC instrument. Each experiment was run in duplicate.

### Phylogenomic analysis

Genbank files (72,556 counts) were downloaded from the NCBI Genbank based on records on the PLSDB database (36, 37). 1,906 files were found to be unannotated and thus removed. Protein BLAST databases were constructed from the remaining 70,650 files. Protein sequences from RK2 KorB (residues 39-149), KorA (full length) and TrbA (full length) were used as queries in protein BLAST searches to identify the corresponding protein in the proteomes. 8,793 proteins were identified and filtered by length: KorB 240-700 residues (5,871 counts), KorA 80-180 residues (1,385 counts) and TrbA 80-460 residues (1,036 counts). The length-filtered collection consisted of 8,292 proteins from 6,736 accessions. Proteins were aligned to create three multiple sequences alignments around the positions F259 of KorB, Y84 of KorA and F104 of TrbA. This collection was dereplicated using mmseqs2 (38) with a cutoff of 90% nucleotide identity resulting in 6,161 unique accessions. The results were analyzed in Excel and plotted using SankeyMATIC (39).

Whole-plasmid similarity was assessed using Average Nucleotide Identity (ANI) calculated via the ANIb method within the pyANI-plus framework (v 1.0.0) (40). The genomes were fragmented into 1020 bp windows, and pairwise alignments were performed using NCBI BLAST+ (v 2.17.0). Pairwise ANI scores, alignment lengths and query coverage were subsequently stored in a SQL database to ensure reproducibility. Plasmids were clustered using a single-linkage method with the classify module of pyANI-plus, which identified clades based on the highest shared identity between any two members of a group. Overall, whole-plasmid ANI analysis showed that plasmids in all subsets (Fig. 5) are generally diverse and partitioned into multiple distinct lineages rather than representing an expansion of a single highly similar backbone.

### Statistics and reproducibility

Experiments were performed in triplicates unless stated otherwise. No statistical method was used to pre-determine the sample size. No data were excluded from the analysis. Details of statistical tests and the *P* values are reported in the legends of relevant figures.

## ACKNOWLEDGEMENTS

We thank Prof. C. M. Thomas for providing *xylE* reporter plasmids, α-KorA and α-TrbA antibodies, and advice during this work. This work is supported by the Lister Institute fellowship (T.B.K.L), by the Wellcome Trust Investigator grant 221776/Z/2/Z (T.B.K.L) that funds T.C.M, by BBSRC Research Grant (APP76057), and by the BBSRC funded Institute Strategic Program Harnessing Biosynthesis for Sustainable Food and Health (HBio) (BB/X01097X/1).

---

**Supplementary Fig. 1.**
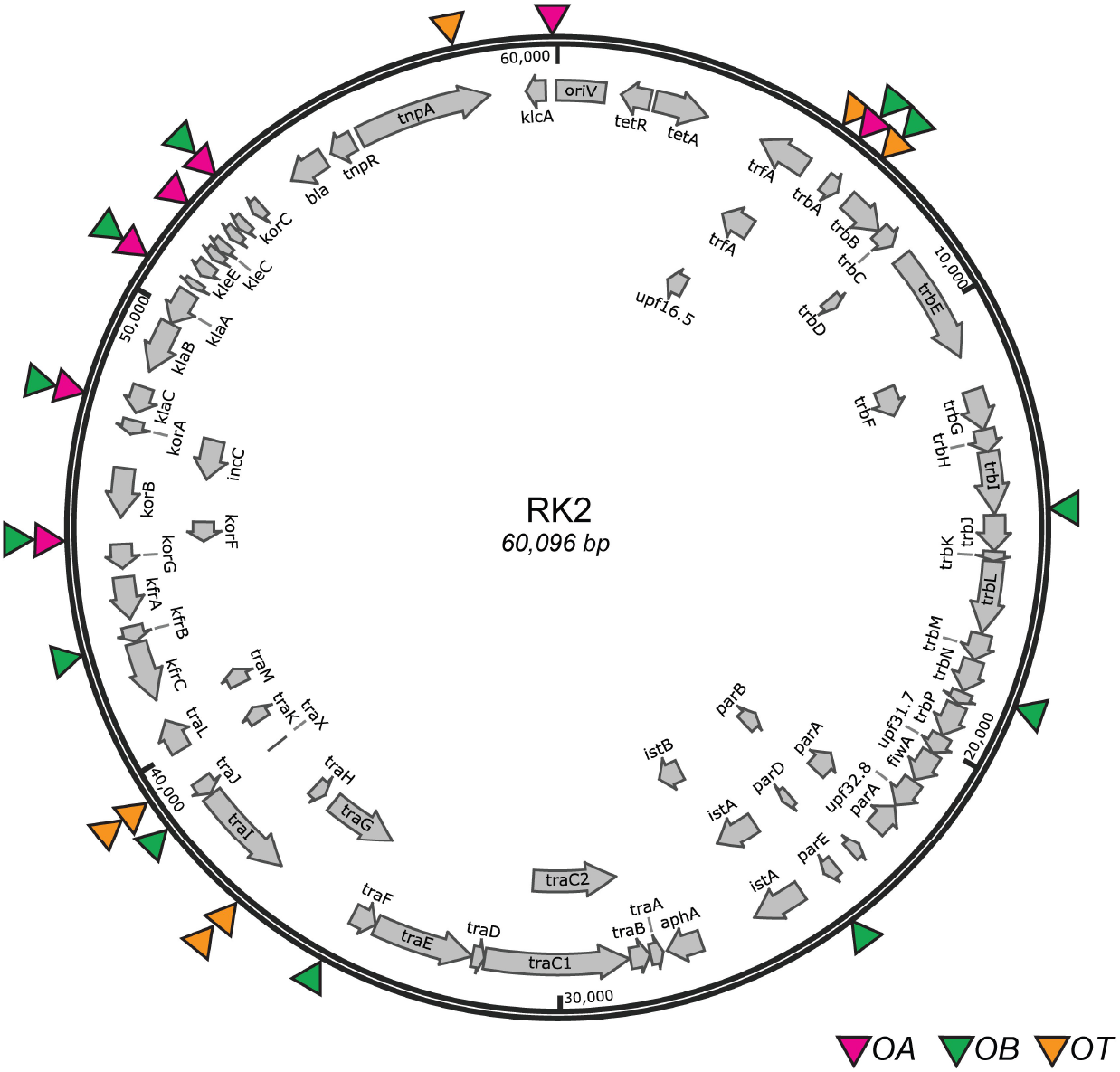
Schematic diagram of RK2 plasmid and its encoded genes. Binding sites of KorB (*OB*, green arrows), of KorA (*OA*, magenta), and of TrbA (*OT*, orange) are also shown on the RK2 plasmid map.

**Supplementary Fig. 2.**
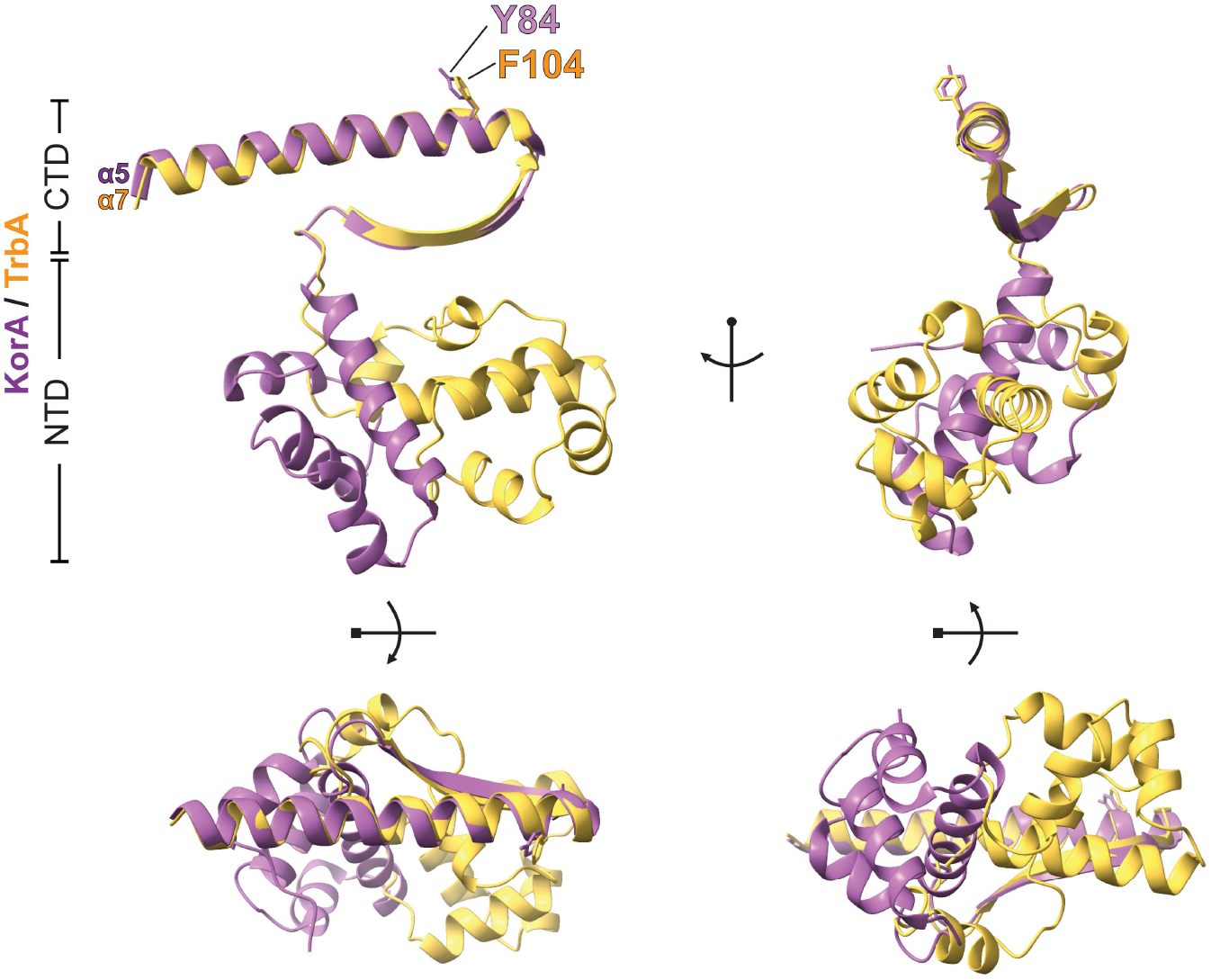
KorA and TrbA share structural homology at their C-terminal domains. Overlay of the AlphaFold2-predicted structure of TrbA monomer (yellow) on KorA monomer (purple). The C-terminal domains of KorA and TrbA closely aligned, while their N-terminal domains do not. The structure of KorA monomer is from the co-crystal structure of a KorBΔN30ΔCTD-KorA-DNA complex (PDB: 8QA9), KorBΔN30ΔCTD and the DNA were omitted for simplicity. Helix α5 (purple) of KorA and helix α7 (yellow) of TrbA, as well as the critical aromatic residue KorA Y84 and TrbA F104, are shown.

**Supplementary Fig. 3.**
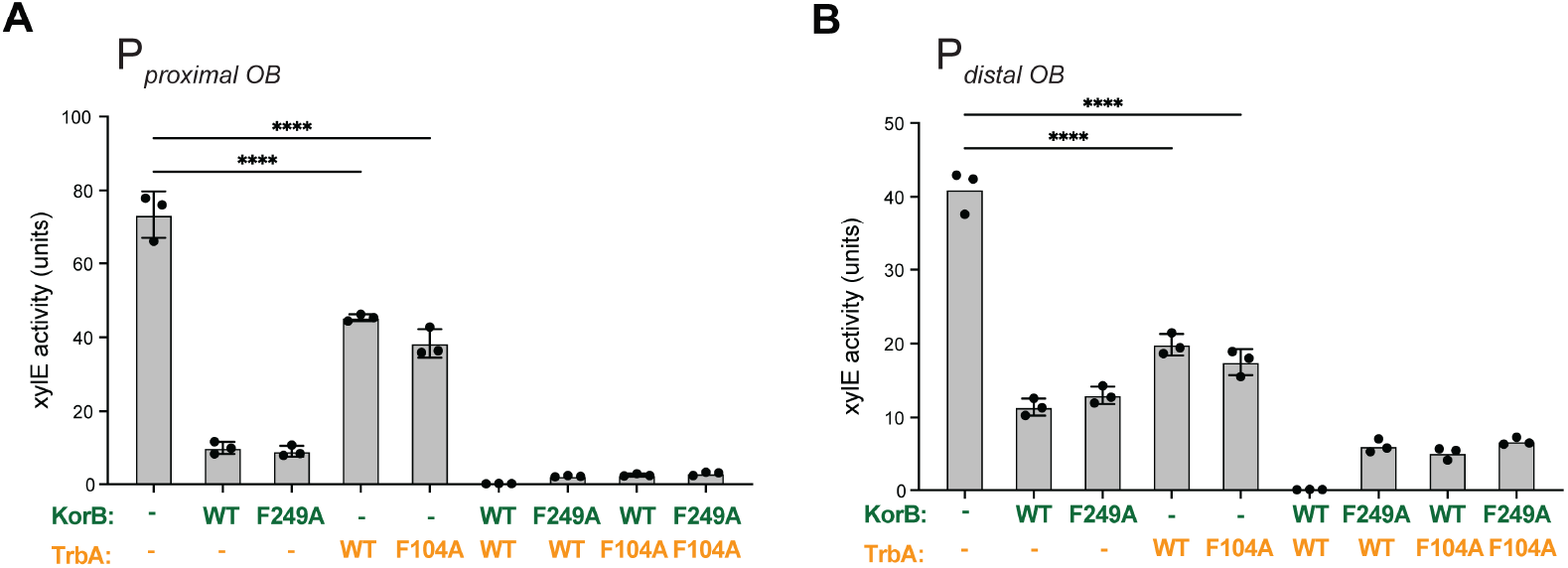
TrbA cooperates with KorB to repress transcription from a distance *in vivo*. Values shown are absolute values of XylE activities from the corresponding panel in Fig. 4B. Experiments were performed in triplicate and mean +/- SDs were presented.

## Notes

### Competing Interest Statement

The authors have declared no competing interest.

### Summary of Updates

Supplementary Tables have now been added to this revision.

